# Systemic mutational rescue in *Escherichia coli* elicited by a valency dependent, high affinity protein DNA interaction

**DOI:** 10.1101/2020.03.09.983320

**Authors:** P.J. Hurd, A. M. Al-Swailem, A.A.A. Bin Dukhyil, Q.I. Sheikh, A. A. Al-Ghanim, L. Alfageih, M. Matin, Yueh Ting Lu, A. Abdalgelel, J. Florence, Mohammed Al-Shemirti, S. Al Harbi, P.E. Brown, D.P. Hornby

**Affiliations:** School of Biological and Chemical Sciences, Queen Mary University of London, London, E1 4NS, UK.; Chief executive officer Saudi Authority for Intellectual Property6531 Al Okaya Rd, Al Sahafah P. O. Box 13321 Riyadh 3059 Saudi Arabia. Email; Medical Laboratories Sciences Department, College of Applied Medical Sciences, Majmaah University, P.O. Box 25529, Riyadh 11476, Saudi Arabia Email; The Krebs Institute, Department of Molecular Biology and Biotechnology, University of Sheffield, Western Bank, Sheffield, S10 2TN; Medical Laboratories Sciences Department, College of Applied Medical Sciences, Qassim University. P.O. Box 6699, Qassim 51452, Saudi Arabia; Department of Biology, Faculty of Sciences,Ferdowsi University of Mashhad, Iran.; Faculty of Science, The University of Kufa, Najaf, Iraq email; Department of Medical Laboratory Science, College of Applied Medical Sciences Al-Quwayiyah, Shaqra University, Kingdom of Saudi Arabi; The Krebs Institute, Department of Molecular Biology and Biotechnology, University of Sheffield, Western Bank, Sheffield, S10 2TN.

**Keywords:** Mutation, rescue, DNA methyltransferase, deletion, insertion, (indel), transposon - mutagenesis, genome engineering, antimicrobial resistance

## Abstract

The controlled formation of sequence specific DNA protein complexes is a fundamental feature of most genetic transactions. Studies of the impact of point mutations on the function of individual components, such as repressors, remains a key aspect of many Systems and Synthetic Biology research programmes. One of the most dramatic systemic consequences of a point mutation is exhibited by the monomeric DNA methyltransferases M.HhaI and M.EcoRII, where substitution of a single, catalytic cysteine by either glycine or alanine, creates a lethal gain of function phenotype.

*In vivo* expression of these point mutants promotes the deposition of high affinity nucleoprotein complexes that arrest replication *in vivo*, causing cell death. Interestingly, it appears that a systemic response to expression of these mutant enzymes is dramatically enhanced when they are expressed as synthetic dimers. A previously unreported form of “mutational rescue” appears to be triggered as a result of networked crosslinking of host cell DNA resulting from an increased valency of nucleoprotein complex formation. This finding may have significance for developing molecular interventions for the controlled regulation of genome function and for the development of novel antimicrobial and anticancer strategies.

## Introduction

The origins of contemporary molecular medicine date back to the pioneering work of Linus Pauling and colleagues (1949) and the subsequent demonstration by Vernon Ingram(1956) that sickle cell anaemia can result from a single missense mutation in the β globingene. However, while the outcome of mutational events is often disadvantageous to an organism, genome stability and mutation are generally considered as two sides of the same coin. On the one hand, the stable and faithful inheritance of genetic information is essential for the viability of a species, on the other, a measured level of change drives evolution at the genetic level. In the simplest examples of genetic mutation, the replication machinery provides the primary “engine” that balances the conflicting demands of stability and change. Our understanding of DNA damage, repair and general genome maintenance has revealed that changes to the genome are brought about by an assortment of mechanisms that range from individual nucleotide events to those involving a form of genetic recombination (Friedberg et al 1995). For example, transposition and deletion events can lead to genome remodelling, promoting both positive and negative outcomes. Furthermore, all organisms encode DNA modifying enzymes, whose activity can impart an additional level of influence on both genome integrity and phenotype. Ongoing research into these phenomena at a systems level and the therapeutic opportunities that may emerge, remain a high priority.

Just as the *lac* operon has proven to be an enduring work-horse for both molecular biology (Jacob and Monod, 1961) and more recently systems biology (Chure et al, 2019), the cytosine-specific C-5 DNA methyltransferases (C5-DNMTs) represent a well-studied class of DNA modifying enzymes, most commonly associated with restriction and modification in prokaryotes, but also as facilitators of a range of epigenetic phenomena (Edwards et al, 2017). *In vivo*, these prokaryotic monomeric enzymes recognise a short DNA sequence (typically between 2-8 base pairs) and methylate one strand of a hemi-methylated DNA duplex, by flipping out the unmethylated, target base (reviewed in Hong & Cheng, 2017). Following S-adenosyl-L-methionine dependent methylation, the target base is returned to the DNA duplex and the enzyme subsequently dissociates. A small number of heterodimeric C5-DNMTs have been described; M.*Aqu*I (Karreman et al, 1986) and M.*Eco*HK31I (Lee et al, 1995), but importantly all possess a single active site. Most bacterial C5-DNMTs form part of a Type II restriction and modification operon. In eukaryotes however (Bestor, 1988; Malagnac, et al, 1997; Kouzminova & Selker, 2001), and some bacteriophage (Trautner et al, 1996) these enzymes are expressed independently of a cognate restriction endonuclease, and their biological roles are diverse, not always understood (reviewed in Edwards et al, 2017).

Many mutants of this class of enzymes have been described, but one unusual mutant stands out: the substitution of the catalytic Cys residue (generally preceded by Pro), for Gly (referred to here as ProGly). Wyszynsky et al (1992) were the first to report the “toxic” phenotype of the ProGly mutation in M.*Eco*RII, and Mi and Roberts (1993) subsequently demonstrated that the same mutation in M.*Hha*I dramatically reduced the dissociation constant for its cognate DNA sequence. Henderson and Kreuzer (2005) went on to confirm, that a key element of the “toxic” phenotype of this mutation results from compromised DNA replication, induced by the expression of abnormally high affinity protein complexes, which in turn inhibit DNA replication. The effects of the ProGly mutant have been compared to the molecular events that follow the administration of anti-cancer drugs including azacytidine (vidaza) (Arimondo et al 2019) and zebularine (Zhou *et al* 2002) or the fluoroquinolone family of antibiotics (see Bax *et al*, 2017). These compounds promote the irreversible accumulation of nucleoprotein complexes at the sites of DNA recognition, which forms the basis of their application in epigenetic medicine (vidaza) and in the treatment of microbial infections (ciprofloxacin). While the specific modes of action of vidaza and zebularine involves incorporation into genomic DNA, followed by the covalent trapping of C5-DNMTs; the fluoroquinolones prevent the completion of the catalytic cycle required for the segregation of bacterial chromosomes (Bax et al, 2017). In this context, Henderson and Kreuzer (2005) have shown that there are clear differences in the cellular responses arising as consequence of the formation of these two persistent nucleoprotein complexes.

During a comparative investigation of the molecular characteristics of drug induced complexes and those formed by the ProGly mutant enzymes, we constructed a range of recombinant C5-DNMTs exhibiting different sequence specificities. In most cases, the expressed enzymes were fused to the C-terminus of a naturally homo-dimeric Glutathione-S- transferase (McTigue et al, 1995; Henderson, et al 1996). One of the most surprising consequences of expressing the C5-DNMTs in this way, was the apparent neutralisation of the “toxic” effect of the ProGly mutation. We report here that the oligomeric status of a point mutation in a DNA modifying enzyme has a profound impact on the systemic response to its expression *in vivo*. These results may provide new insights into our understanding of the role of genome topology in genome surveillance and maintenance, and point to important considerations in understanding the interplay between the sequence and the operational valency of a protein in a wider evolutionary context.

## Results

### 1. Synthetic oligomerisation directs the host response to high affinity DNA recognition

A range of plasmids encoding the prokaryotic C5-DNMT enzymes, M.HhaI, M.MSPRI and M.MspI were generated and provided the starting point for all genetic constructs in the experiments described here. Their general properties are summarised in Table 1. Comparative transformation efficiency was used to assess the DNA modification proficiency of all plasmids encoded enzymes, using the protocol described by Raleigh and Wilson (1986). As expected, plasmid transformation of *E.coli* strains AB1157 (*mcr*+) and DH5αMCR (*mcr−*) gave the anticipated results with respect to DNA methylation proficiency of each plasmid encoding a monospecific C5-DNMT (see Table 1). The expression of missense mutants where catalytic Cys to Gly (ProGly mutation) is incorporated, significantly reduces host viability, as demonstrated by relative transformation efficiencies shown in Figs 1 a and b. These results are in full agreement with the observations of Wyszinsky et al (1992) and Mi and Roberts (1993). Furthermore, plasmids encoding ProGly mutations yielded negligible numbers of colonies. Superficially, the impact of the ProGly mutation appears similar to the restriction phenomenon observed when appropriately methylated plasmids are eliminated from *mcr*+ strains (Raleigh and Wilson, 1986).

**Table 1.**
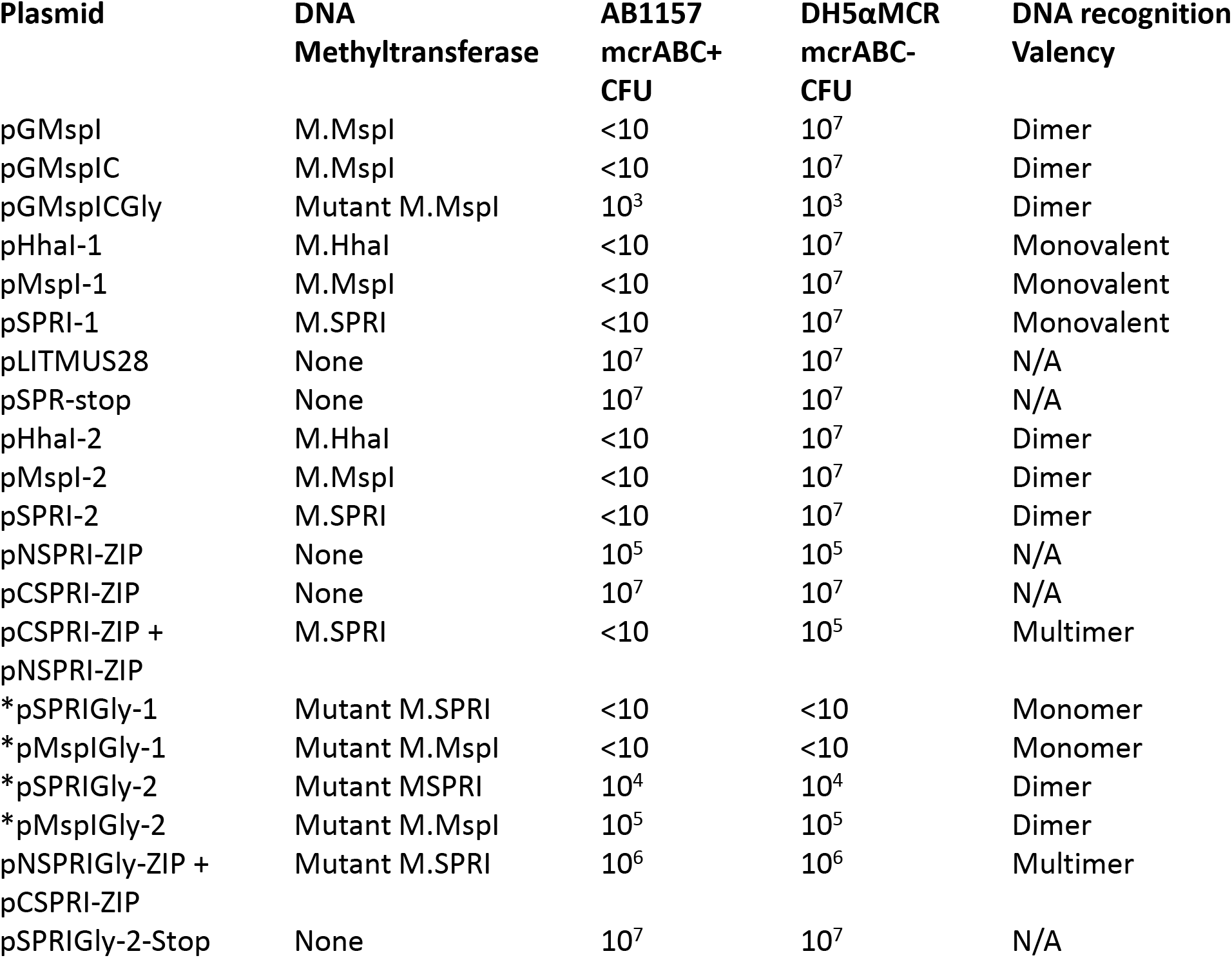
Plasmid constructs used in this work. The relative transformation efficiencies are expressed using pSPR-stop and pLITMUS28 (New England Biolabs) as reference plasmids. *These plasmids were introduced following Kan insertion to inactivate the C5-DNMT during Gly mutant construction and subsequent removal (by restriction and subsequent ligation, as shown in Fig 2b).

**Fig. 1.**
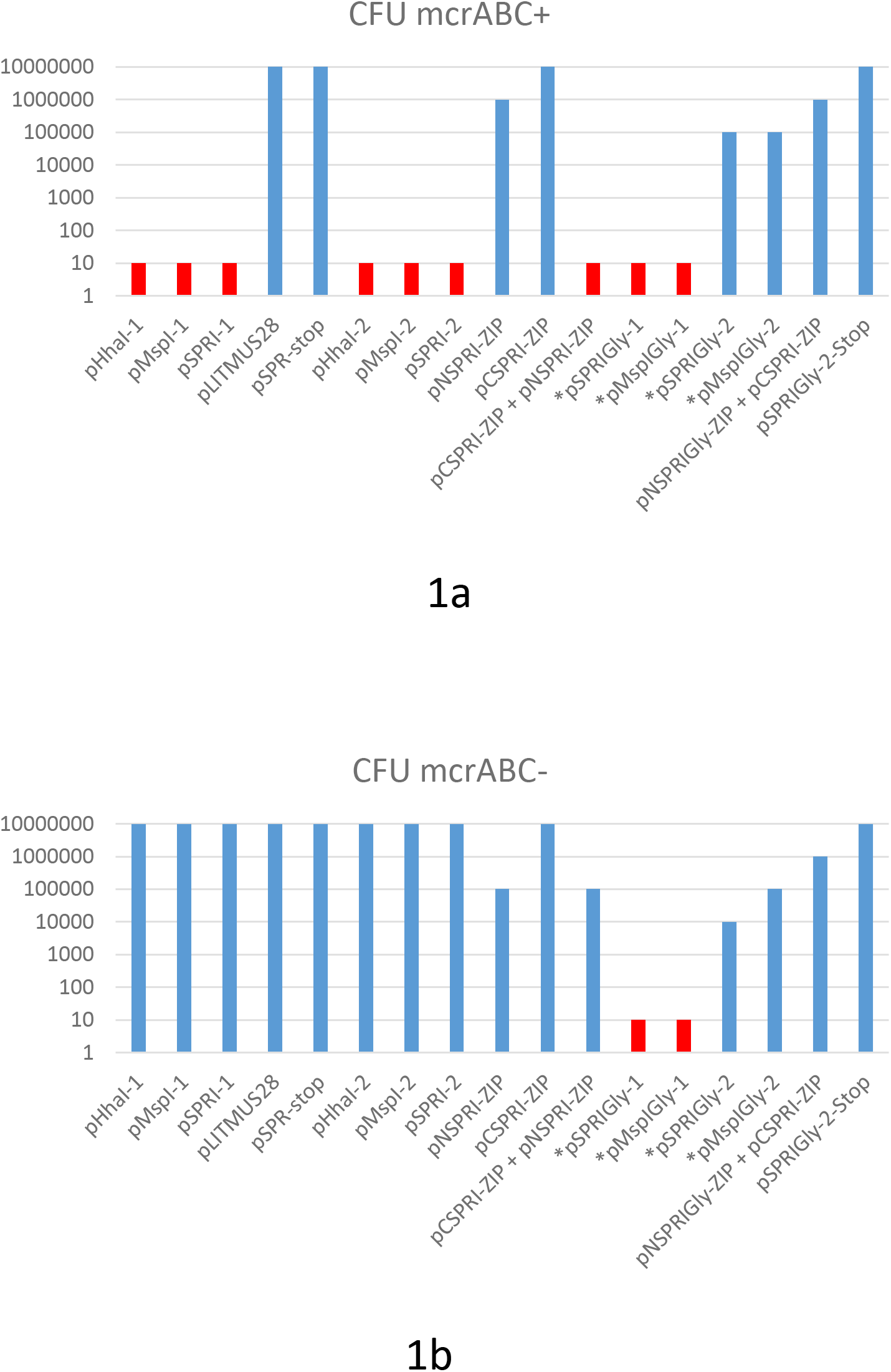
Log plot of the transformation efficiencies of plasmids used in this work. (a) *E.coli* AB1157 (mcrABC+) was the host for the transformation in each case of 10ng of supercoiled plasmid DNA. Plasmids indicated by and asterisk contain a ProGly mutation and were digested with a single restriction endonuclease to excise a kanamycin gene cassette, followed by re-ligation. In these cases, all CFUs reported are ampicillin resistant and kanamycin sensitive. Transformation experiments involving the monomeric ProGly mutant encoding plasmids are coloured red. The same experiments in (a) are shown in (b) with *E.coli* DH5αMRC acting as recipient.

As part of a series of experiments investigating the active site of several bacterial C5-DNMTs, plasmid expression vectors encoding Glutathione-S-transferase (GST) were constructed as described elsewhere for M.MspI (Taylor et al, 1993). The general features of this group of plasmids and their expression products are provided in Table 1. As with the WT monomeric forms of the encoded enzymes, the polypeptides retain DNA methylation proficiency as artificial dimers (a consequence of the obligate dimerization of GST). All plasmids expressing GST fusions of M.MspI, M.SPRI and M.HhaI are fully methylated and are consequently restricted by mcrABC+ strains (eg AB1157), but can be stably maintained in *E.coli* DH5αMCR (Taylor et al, 1993).

Surprisingly, following substitution of the ProCys motif at the active site of M.MspI with ProGly (expressed from pGMspICGly, illustrated in Fig. 2a), and in contrast to the toxicity observed following transformation with plasmids encoding monomeric ProGly mutant C5-DNMTs, transformation efficiency was only moderately reduced (see Table 1 and Fig.1). Subsequent extraction and purification of the GST fusion as described elsewhere (Taylor et al, 1993;), from colonies harbouring the plasmid pGMspICGly revealed an enzymatically inactive GST fusion protein of the expected M_r_, as determined by SDS polyacrylamide gel electrophoresis (data not shown). However, nucleotide sequencing of pGMspICGly revealed a 30bp in-frame deletion (eliminating 10 amino acids from the catalytic domain) in all plasmids recovered from transformed colonies. Similarly, two other ProGly mutant C5-DNMTs also showed only marginal reductions in transformation efficiencies in both AB1157 and DH5αMCR hosts (see Table 1 and Fig.1). In these experiments, the ProGly mutation was introduced into the gene after a first step in which a kanamycin cassette was inserted to disrupt the C5-DNMT gene (as shown in Fig. 2b). The protocol used to remove the kanamycin cassette in this way led to a reduction in transformation efficiencies compared with supercoiled plasmid controls, as would be expected, but proved a more efficient method for reproducible evaluation of transformation efficiencies. A profile of transformation efficiencies of all plasmids used in this work is given in Table 1 and Figure 1. It is clear from these data that the observations of Wyszinsky et al (1992) and Mi and Roberts (1993) only apply to plasmids encoding monomeric ProGly mutant C5-DNMTs. Therefore, synthetic, GST dimers overcome the barrier to transformation observed when plasmids encode monomeric C5-DNMT ProGly mutants. It should be noted that, in view of the incorporation of plasmid mediated ampicillin selection in the experimental design, only cells retaining replication competence and antibiotic resistance are expected to be recovered in these experiments.

**Fig 2a.**
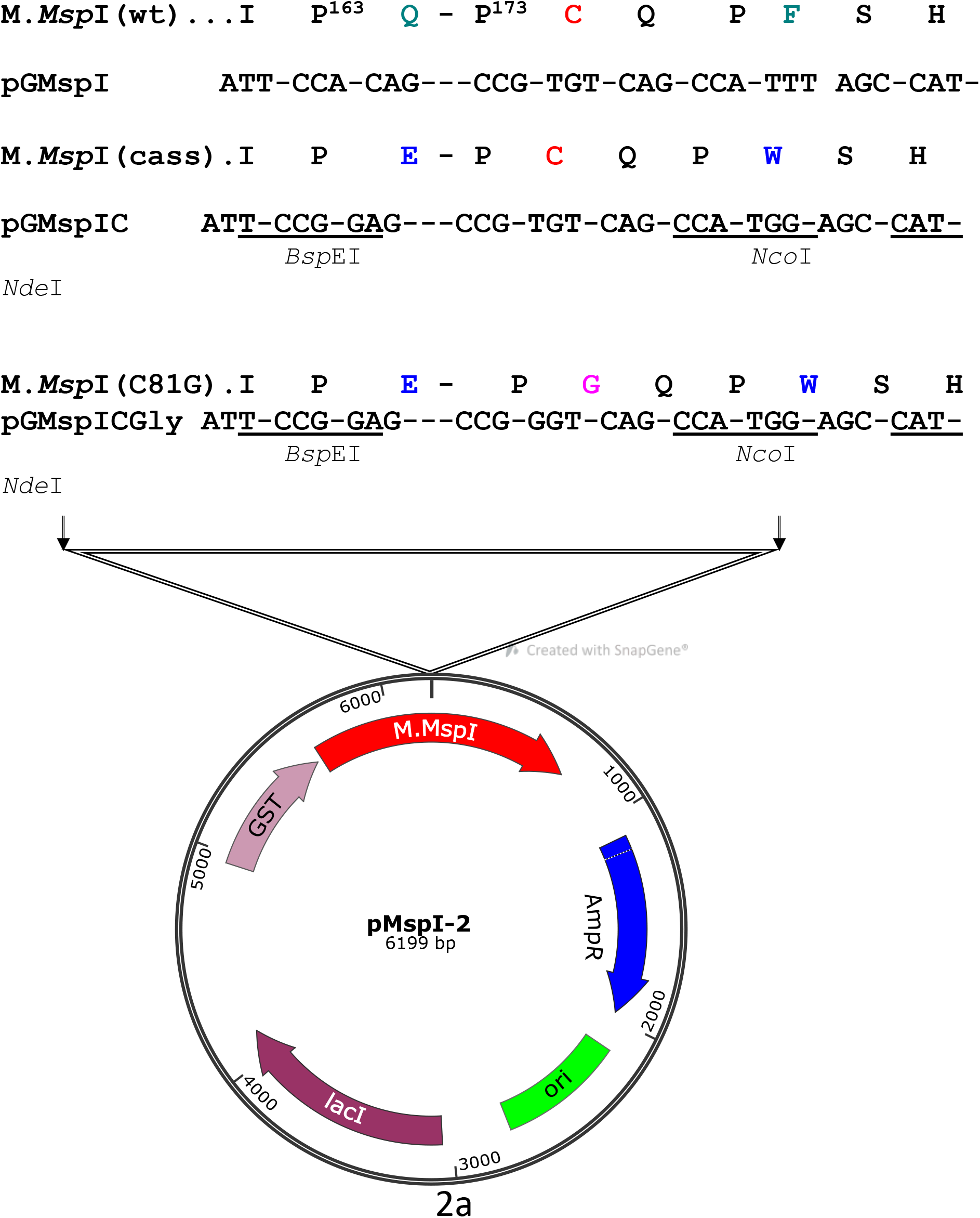
Construction of the ProGly mutant in pMspI-2. The sequence encoding the catalytic loop of M.MspI was modified by oligonucleotide exchange methodology. The resultant plasmid contained unique BspEI and NcoI sites, facilitating the codon change as shown in a second restriction and re-ligation step.

**Fig 2b.**
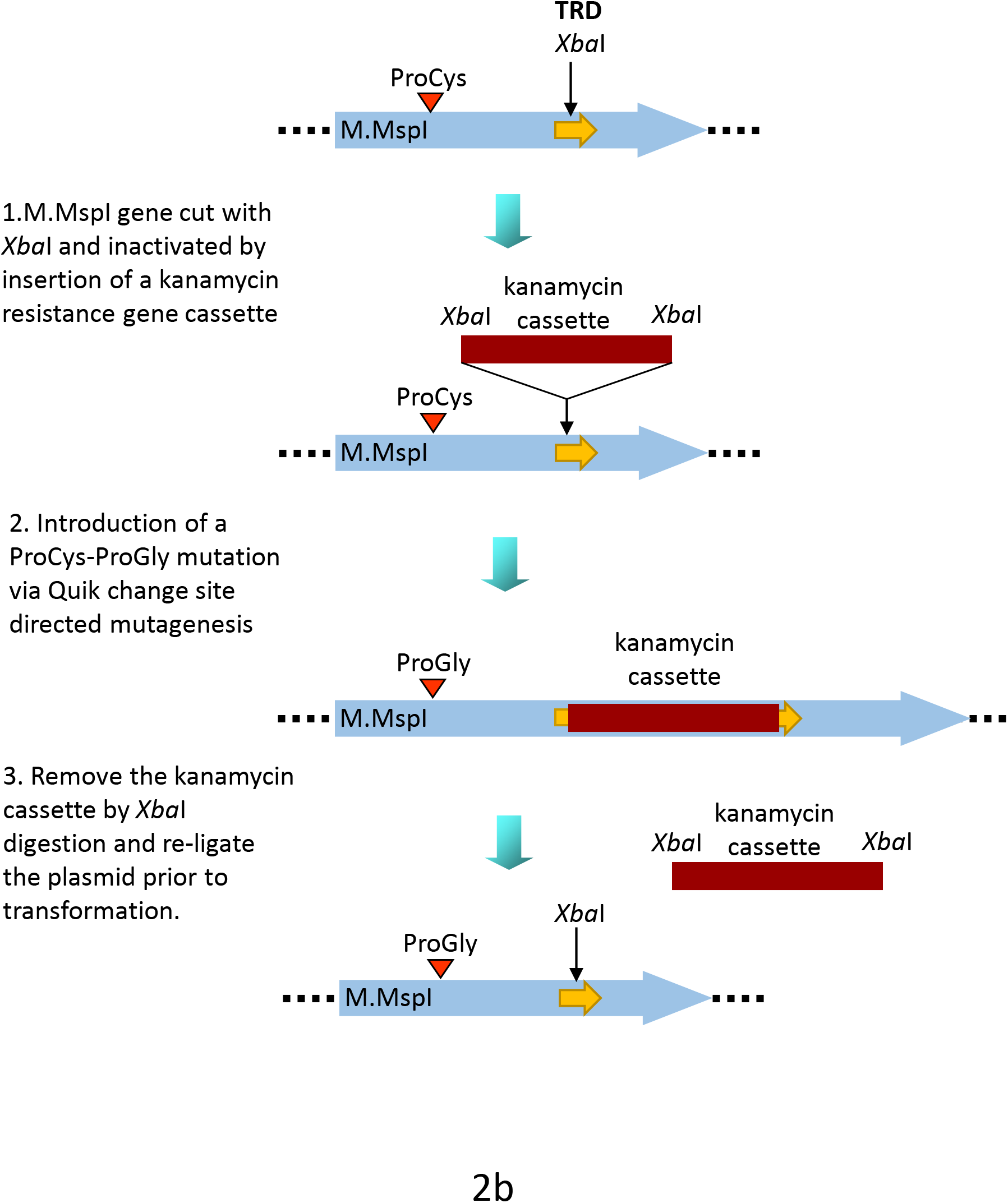
Alternative methodology for the generation of the ProGly mutants. In this case the plasmid encodes a GST fusion of M.MspI. An XbaI gene cassette encoding the kanamycin resistance gene from pUC4K (GE Healthcare) was inserted into the unique XbaI restriction site in the region encoding the Target Recognition Domain (TRD) of the C5-DNMT. Following site directed mutation of the catalytic Cys codon Gly, the plasmid was re-cut with XbaI, re-ligated and transformants selected for ampicillin resistance and kanamycin sensitivity.

Given the sequence specific nature of C5-DNMTs, we reasoned that output plasmids might show patterns of sequence similarity in the vicinity of the mutational rescue process. However, this is not the case and is exemplified by the analysis of a series of mutants recovered from a pSPRIGly-2 transformation experiment. As can be seen in Fig.3, the nucleotide sequences of 5 classes of deletion mutants showed no similarities in sequence at the boundaries of the deletion, and the deletion events are unrelated to the recognition sequences of this multi-specific C5-DNMT (GGCC, CCGG and CCA/TGG). To date, we have not detected any patterns following nucleotide sequencing of over 100 plasmids encoding bivalent ProGly mutant C5-DNMTs.

**Figure 3.**
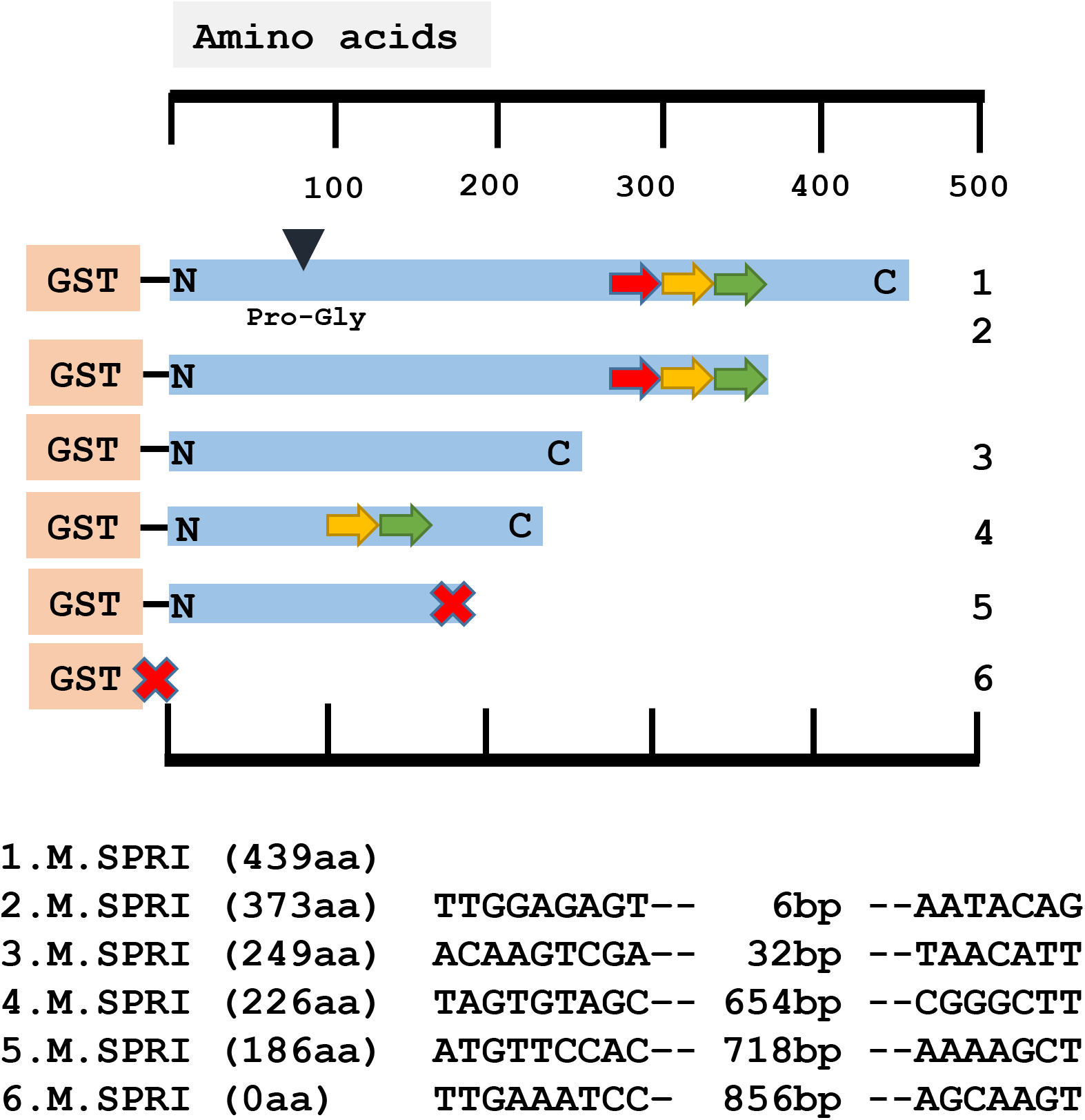
Comparative NUCLEOTIDE sequence analysis of plasmids recovered following transformation of pSPRIGly-2. The coding sequence is indicated by the blue bar, with the N and C termini labelled and the mutation of the catalytic Cys identified by the black triangle as Pro-Gly. The three TRDs are shown as coloured arrows. The sequences at the boundaries of the deletion events are shown below for each plasmid. The large red cross indicates the presence of a premature stop codon.

### 2. The lethality of ProGly mutants is dependent on multivalent expression of C5-DNMTs

In order to further investigate the role of increased valency of high affinity nucleoprotein complex formation, a pair of compatible plasmids were constructed in which the catalytic domain of M.SPRI was expressed independently from one plasmid (pNSPRIGly-ZIP) and the C-terminal DNA target recognition domain (TRD) expressed from a second (pCSPRI-ZIP). Both segments were expressed as GST fusions, and functional assembly of the two separated domains was achieved by tagging the C-terminus of the catalytic domain with the b-ZIP sequence from rat C/EBPα. The same sequence was appended to the TRDregion. One additional advantage of this pair of constructs lay in their ability to reconstitute a functional C5-DNMT or a ProGly mutant via sequential transformation of supercoiled plasmids, thereby eliminating the kanamycin excision step. Interestingly, both transposable and IS element insertions were the hallmark of the loss of C5-DNMT ProGly “toxicity” in the b-ZIP constructs (see Fig. 4). However, while insertion mutants dominated transformation plates, deletions were also recovered. This experiment independently confirms the requirement for multivalent DNA complex formation, since the combinatorial nature of b-ZIP transcription factors is well established (Rodríguez-Martínez et al, 2017). Moreover, because of the promiscuity of b-ZIP motifs, it is likely that a spectrum of possible permutations of oligomers are expressed *in vivo*, including not only monomeric and dimeric forms, but also polyvalent DNA binding proteins. The results of all transformation experiments are summarised schematically in Fig. 5.

**Figure 4.**
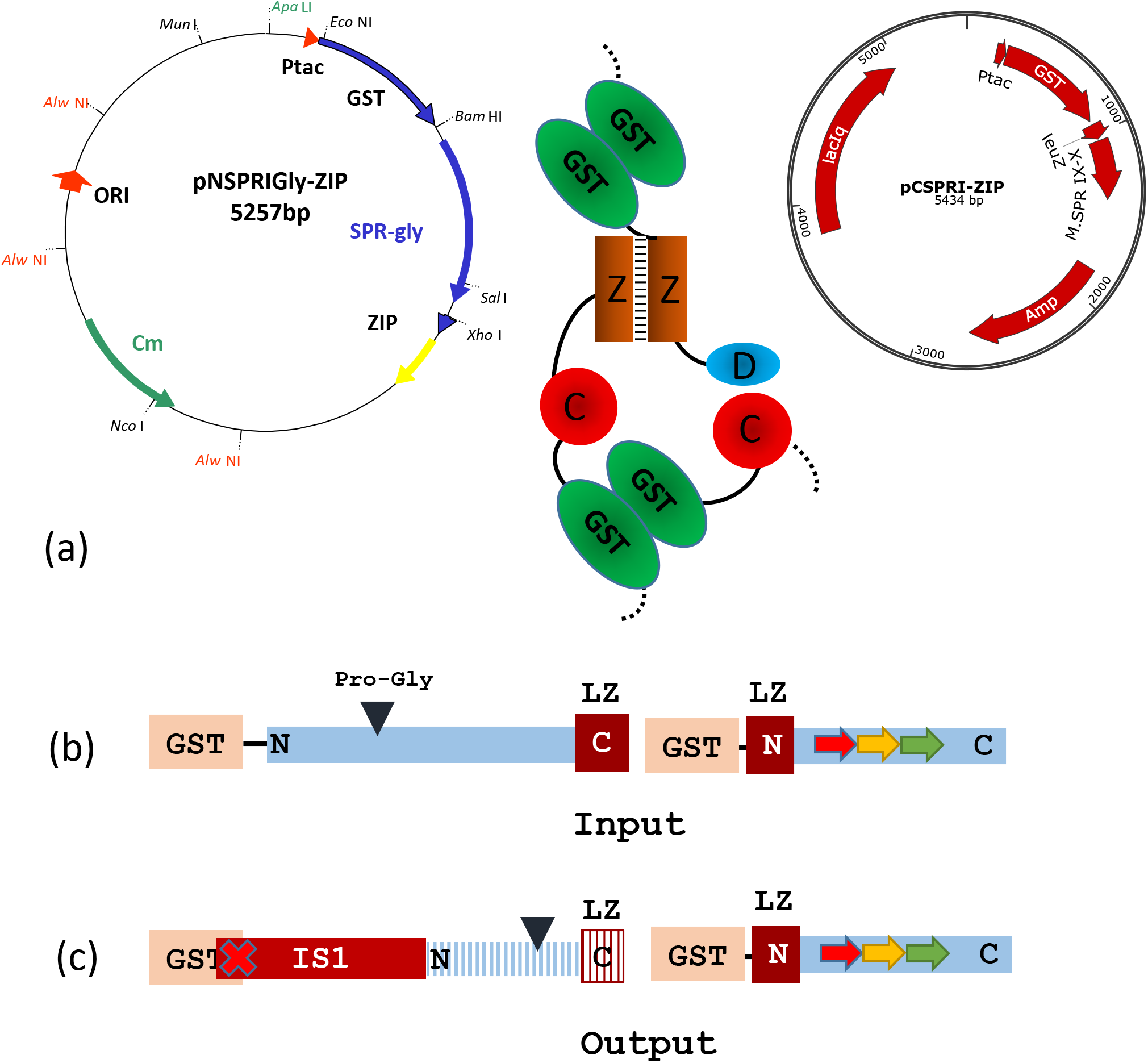
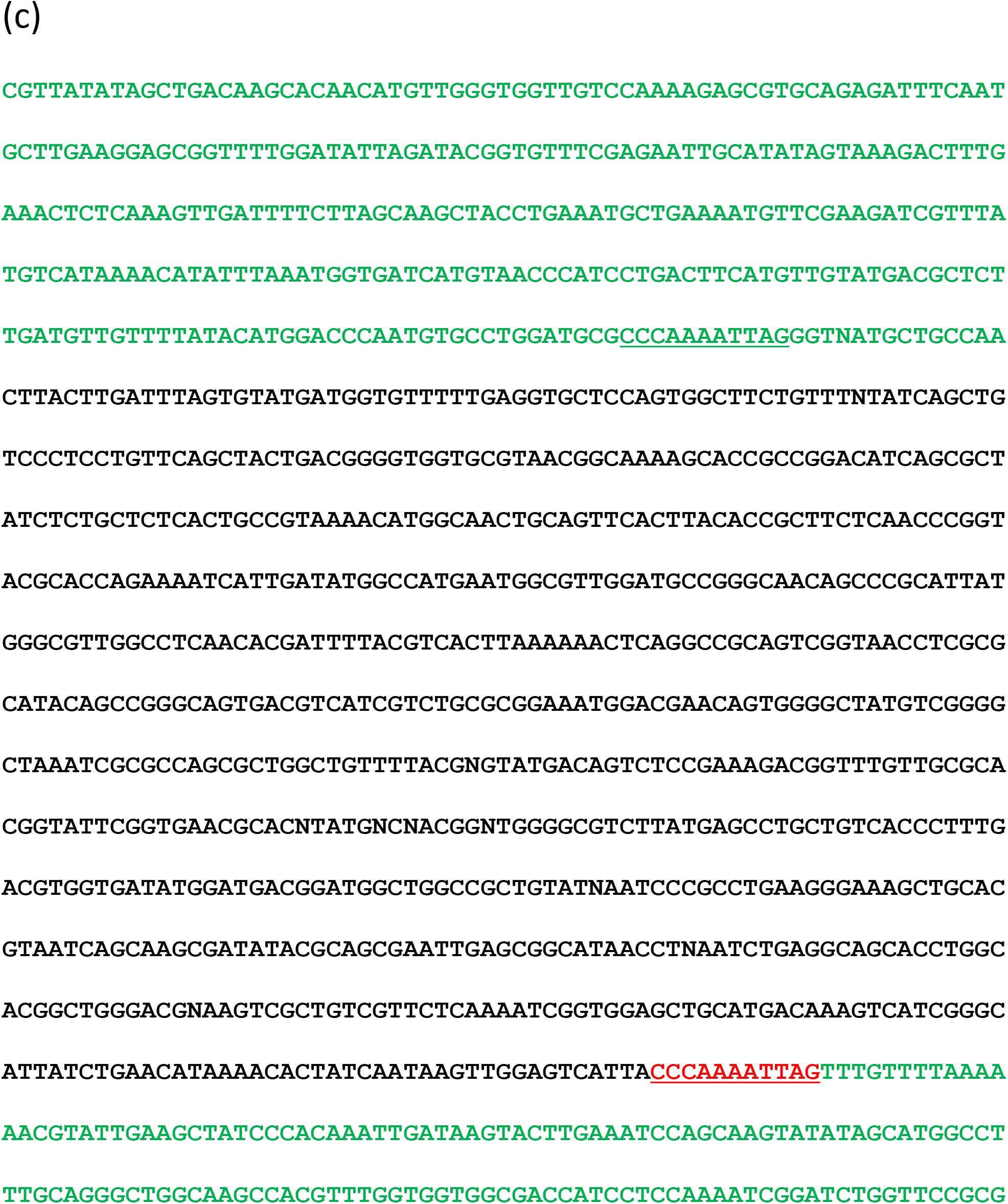
A typical result obtained following transformation of AB1157 with a pair of compatible plasmids designed to generate a heterodimeric M.SPRI enzyme, carrying the ProGly mutation. (a) The assembled enzyme is shown schematically with the b-ZIP motifs enhancing assembly. (b) The schematic diagram shows the input open reading frames expressed from the complementary plasmids. The output ORFs indicate that ISI insertion leads to premature termination of the N-terminal GST catalytic domain (blue stripes), with retention of the C terminal domain. (c) The nucleotide sequence in the vicinity of the insertion event.

**Fig 5a.**
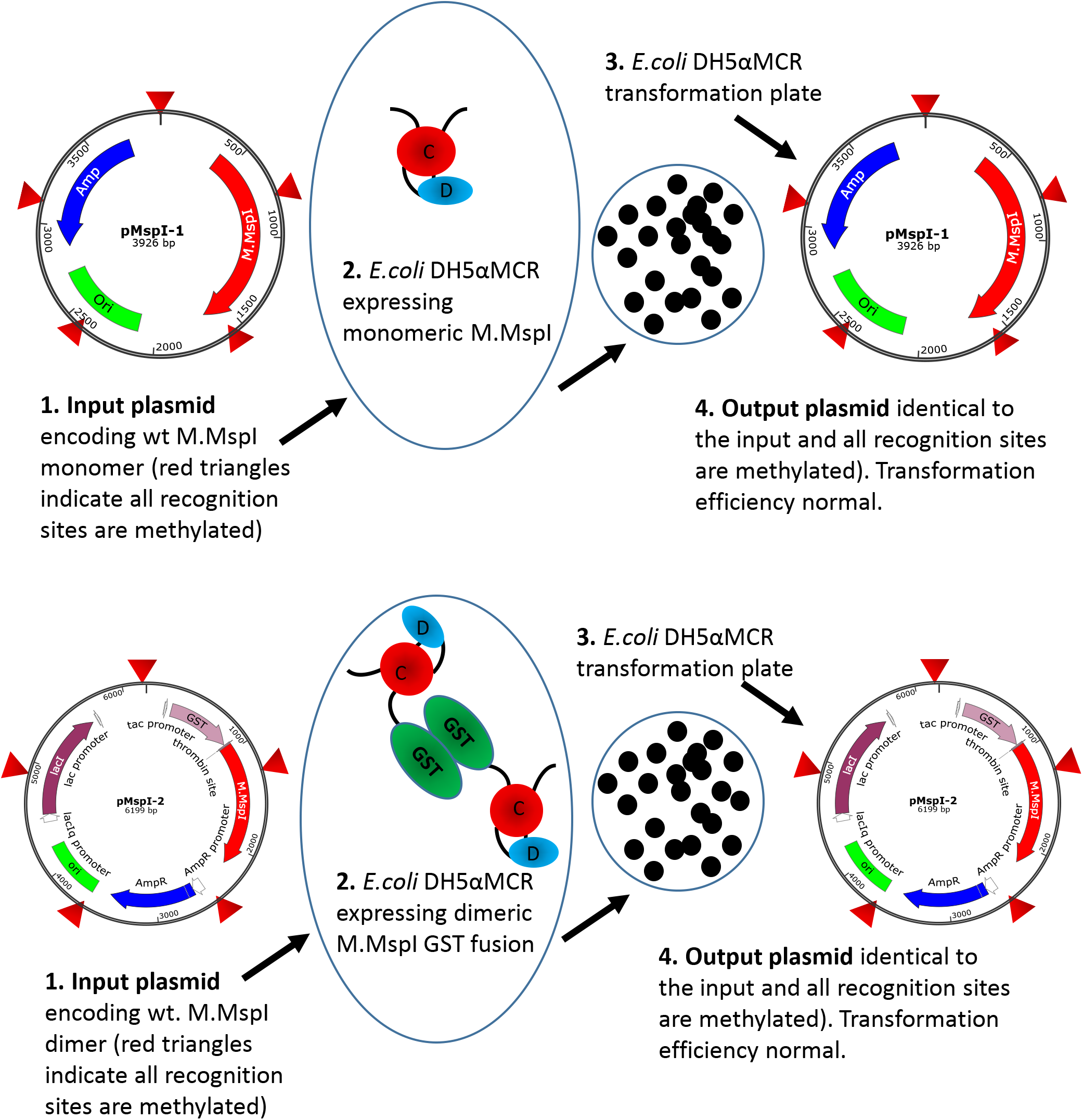
Schematic representation of the transformation experiments exemplified with M.MspI. The input plasmids both lead to the expression of methylation proficient enzymes. In the case of pMspI-1 the enzyme is monomeric whereas the enzyme expressed from pMspI-2 is a GST fusion and therefore dimeric. Transformation efficiencies are comparable and the output plasmids are unchanged.

**Fig. 5b.**
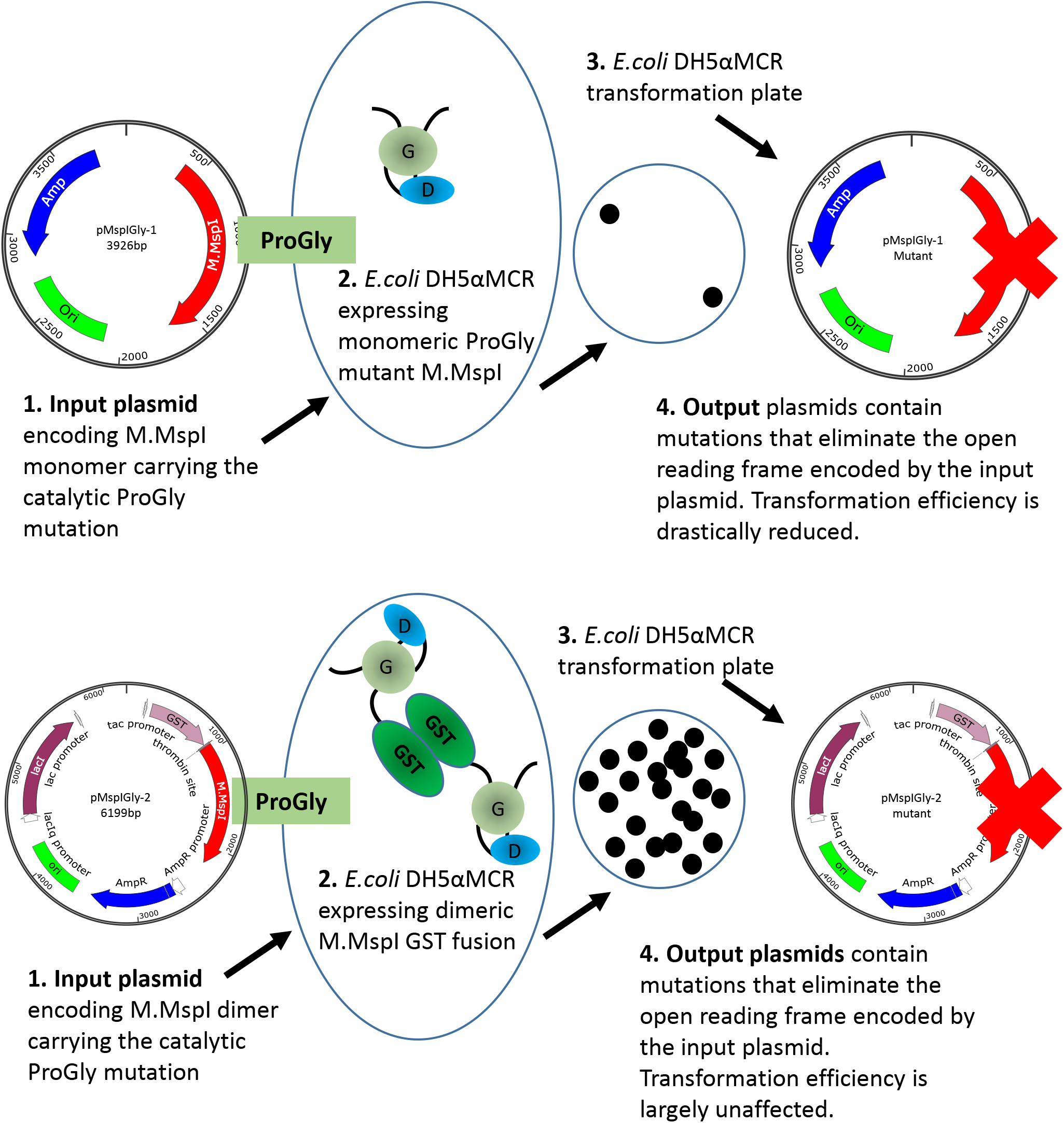
The diagram illustrates the different consequences arising from the transformation of plasmids encoding the monomeric ProGly mutant in the enzyme M.MspI (pMspIGly-1) and the dimeric form (pMspIGly-2). In both cases output plasmids differ from the input plasmids, but importantly, the transformation efficiencies observed when the ProGly mutant is presented *in vivo* in a bivalent form are comparable with those observed using wt plasmids. (The bold red cross indicates that plasmids carry either, deletion, insertion, missense or nonsense mutations. Revertants in which the ProGly sequence is altered to ProCys would be viable, but have not been recovered to date.)

### 3. Mutational rescue is restricted to plasmids and not the genome

In order to establish whether the enhanced levels of mutation induced by the dimeric ProGly mutants, is restricted to plasmid DNA, several colonies were subjected to whole genome sequencing. In this experiment, genomic DNA from *E.coli* DH5αMCR carrying a deletion of the *mcrBC* locus was analysed alongside that from AB1157 strains transformed with pSPRIGly-2, with and without the kanamycin gene cassette. Genome sequencing was carried out by MicrobesNG at the University of Birmingham UK and sequences compiled and analysed using Integrative Genomics Viewer Software (version 2.6.3). Alignment of the genome sequences showed no evidence for genome wide indel events (see Fig. 6). The presence of the *mcrBC* deletion in strain DH5αMCR acted as a reference point for genome comparisons. BLAST was used to identify the pSPRIGly-2 plasmid in each genome assembly: a search was carried out on the partially assembled genome files, using the pSPRIGly-2 plasmid file as the query. The resultant matches identified in which nodes the plasmid could be located, in each of the genome sequences. From this, the plasmids could be identified as encoding either the ProGly mutant, or the kanamycin insertion construct which had not been primed by removal of the insertion sequence. All plasmid sequences revealed inactivating mutations as seen in the earlier plasmid isolation and sequencing experiments. In addition, the kanamycin sequence is clearly present in transformants that were not processed in advance of transformation.

**Figure 6.**
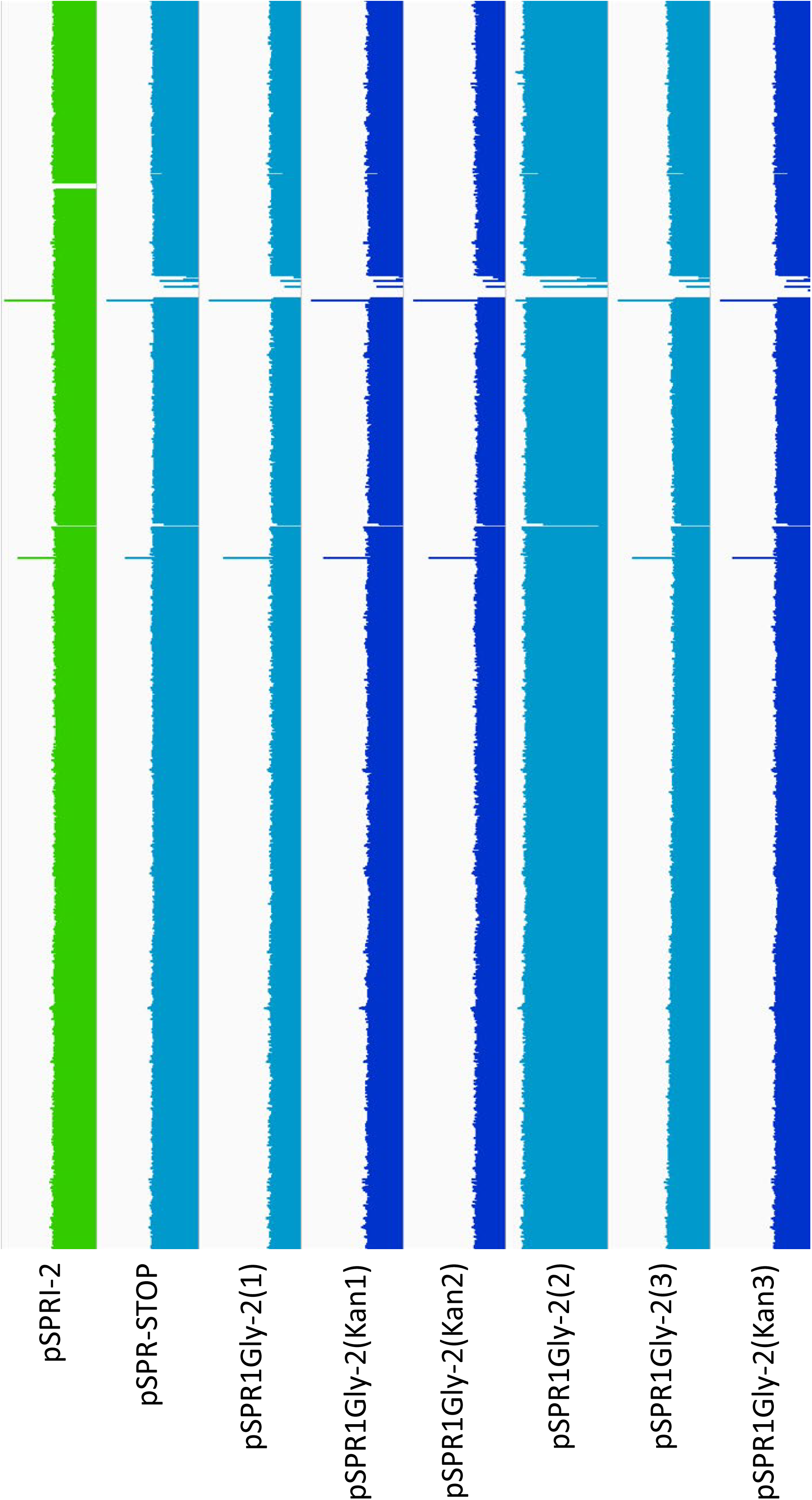
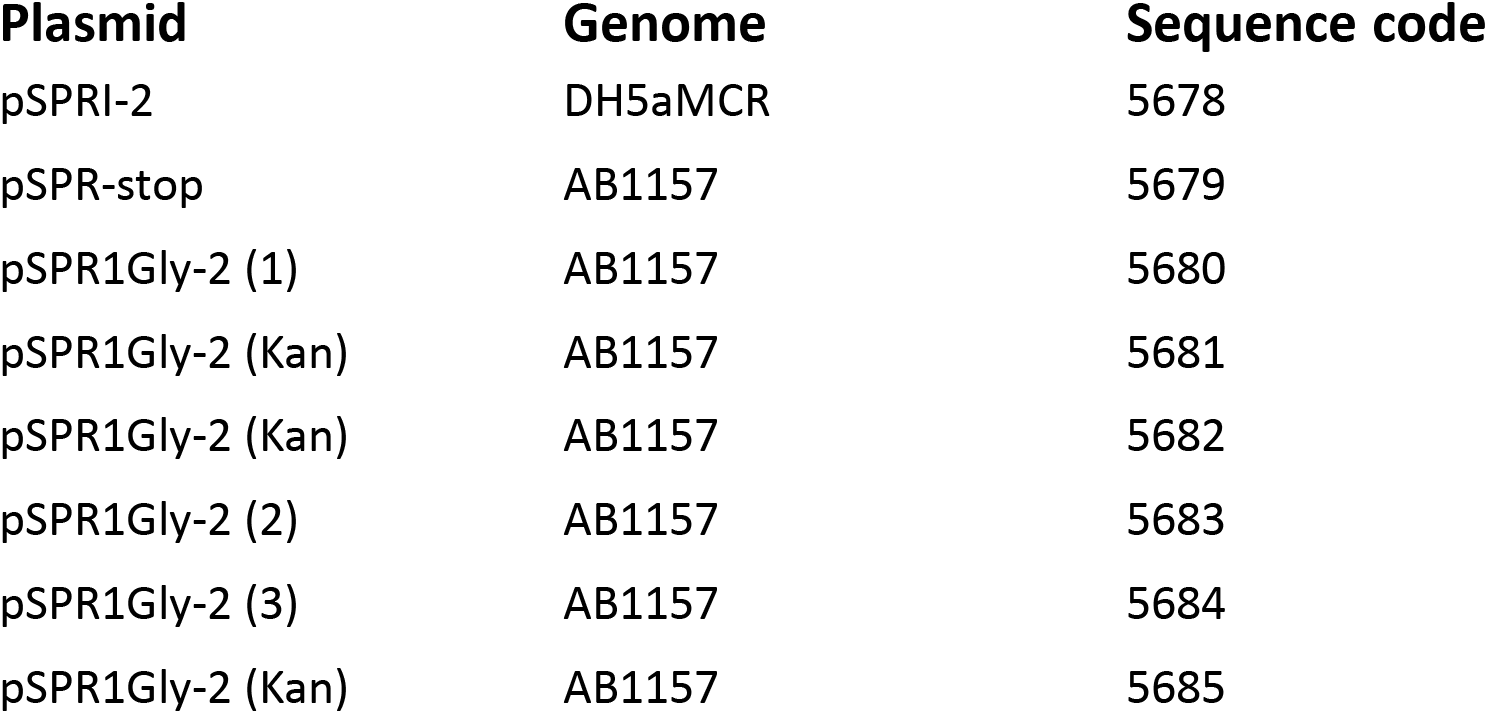
Comparative analysis of genomic DNA following transformation with ProGly mutants expressed from the plasmid pSPRI-2, with or without excision of the kanamycin cassette as indicated. Genome sequences were constructed using the *E. coli* DH1 strain (NC_017625) genome as a template. The number of reads at each point of the genome were plotted using IGV software and compared. Because some regions of the genome were well covered, while others were not, a logarithmic scale was used.

A further comparative BLAST analysis of 50 *E.coli* genes showed no sequence variation across all experimental strains. This result is not unexpected, given that the cells recovered retain viability under ampicillin selection as judged by standard overnight growth protocols. An additional evaluation of the location of transposable elements was carried out, and in this respect there appears to be no difference between the genomes sequenced. Through the comparison of these genomes, it is clear that there are no large rearrangements, deletions or insertions present in strains that have recovered from expression of a bivalent ProGly mutant C5-DNMT.

### 4. Mutants that abrogate mutational rescue

The catalytic pathway of all C5-DNMTs is known to involve flipping of the target cytosine from the DNA helix into the catalytic site, followed by the formation of a transient covalent complex. The difficulties in isolating sufficient quantities of ProGly mutants, have made structure determination an insurmountable challenge to date. However, it is not unreasonable to speculate that the unusually high affinity for DNA exhibited by ProGly mutants, results from the formation of an irreversible base flipped complex. The determinants of sequence specific base flipping reside in the C-terminal domain of C5-DNMTs and comprise the TRD and conserved motifs IX and X (Lauster et al 1989; Pósfai et al, 1989; Klimasauskas et al 1991). In order to investigate the possibility that the ProGly mutants show high affinity recognition through an interplay between the catalytic domain and the TRD domain, a series of M.HhaI mutants were generated using error-prone PCR (Biles and Connolly, 2004) and the library recombined with a plasmid (pHhaI-2) carrying the ProGly mutation. Colonies recovered from this “library” were initially screened for plasmid integrity and were subsequently sequenced. Two classes of plasmids were obtained: those retaining the characteristics of the input plasmid while neutralizing the ProGly effect, and those modified by the mutational rescue phenomenon described above. Inspection of the former class of plasmids revealed that the secondary mutations found in the sequence encoding the C-terminal domain were associated with amino acids known to play a role in the stabilisation of the flipped cytosine ring or in stabilising the structure of the entire domain. Several of these mutants are shown in the context of the secondary structure of M.HhaI in Fig. 7: there is a clustering of mutations around the structural elements associated with stabilisation of the flipped base and also in the C-terminal, conserved structural elements that in mono-specific DNMTs interact with the TRD and confer sequence specificity on the enzyme (Klimasauskas et al, 1994r).

**Fig. 7.**
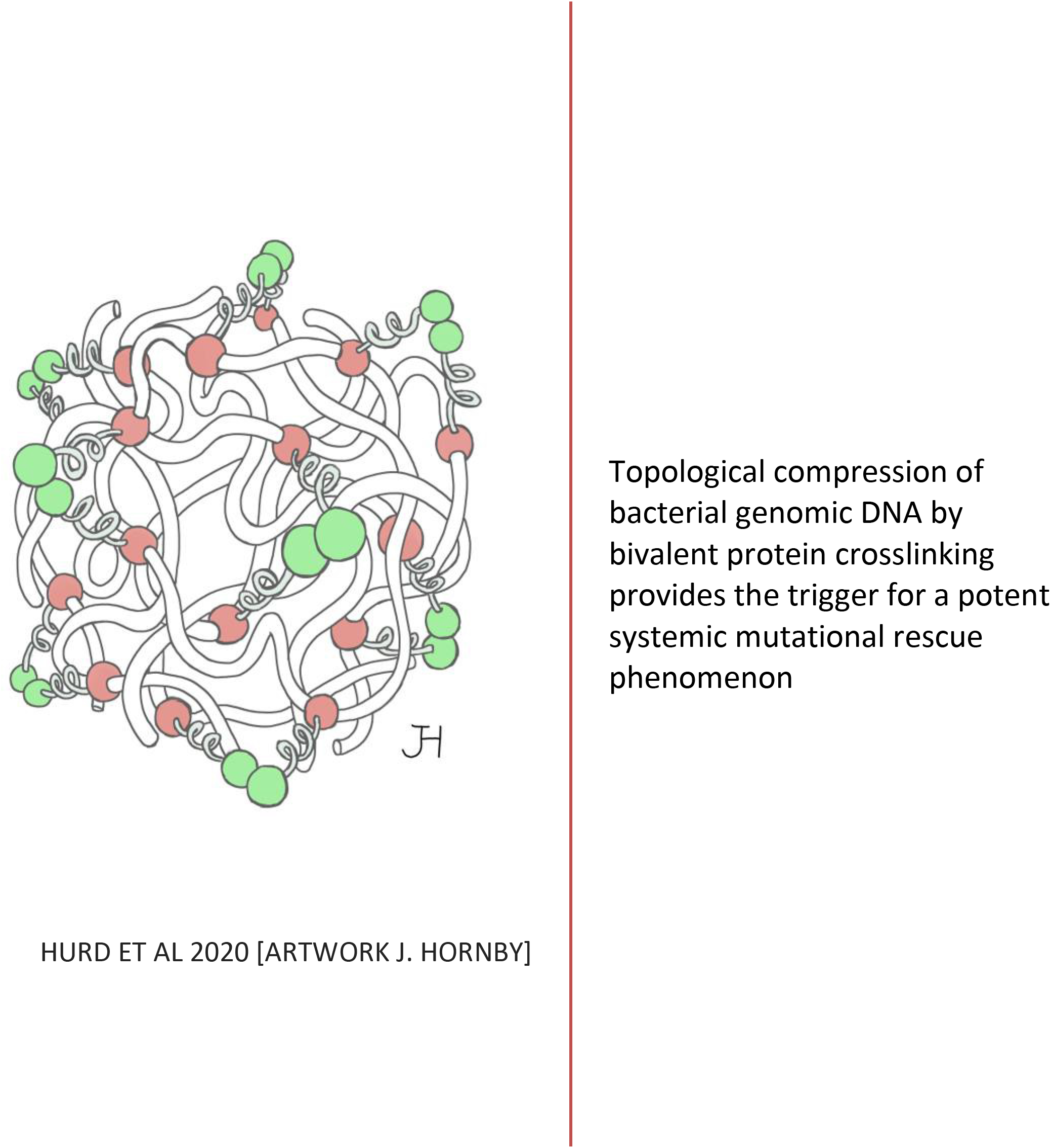
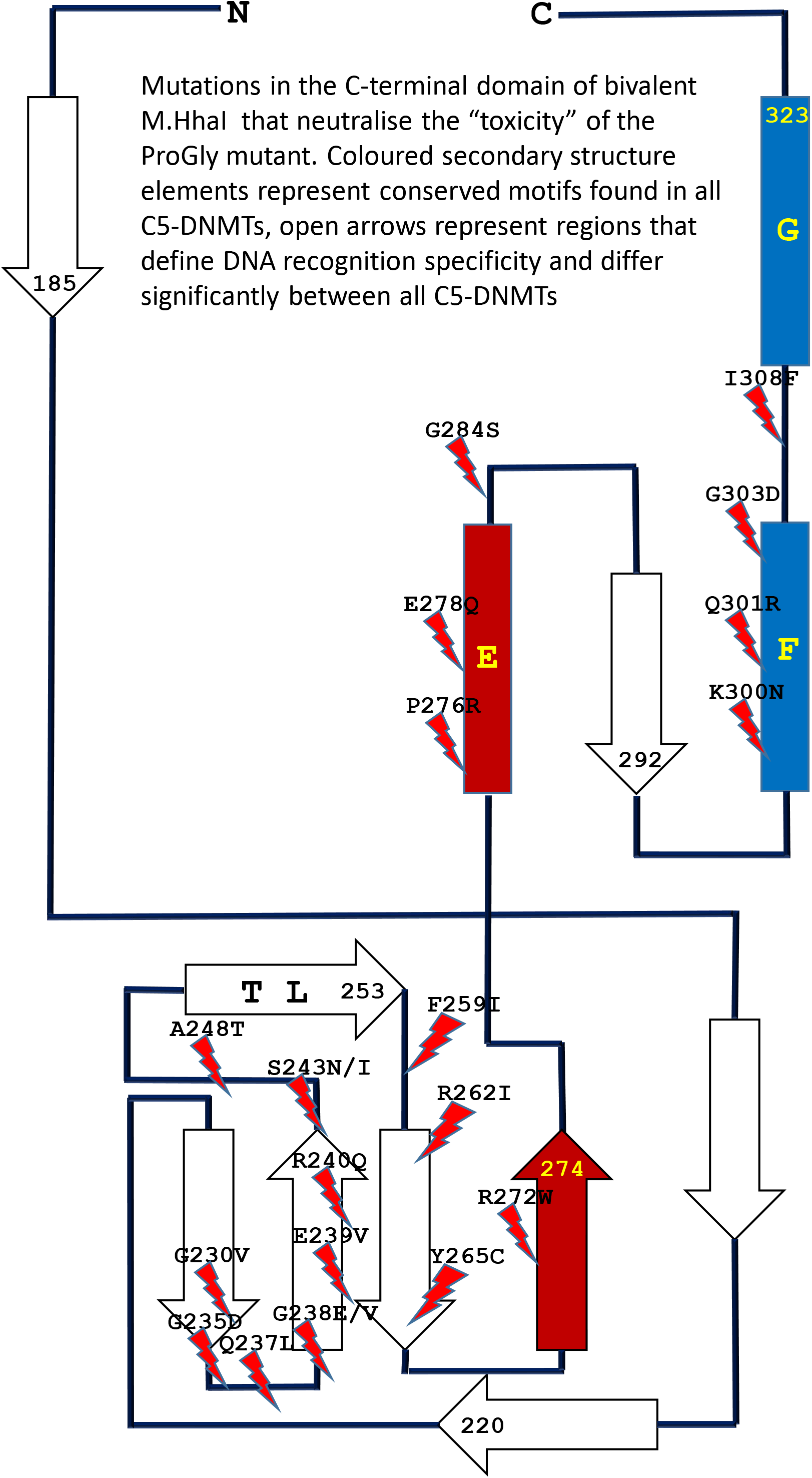
Schematic diagram showing the locations of point mutations recovered from cells in which the sequence encoding the catalytic N-terminal domain of an M.HhaI ProGly mutant, expressed as a GST fusion, is combined with a library of random mutants obtained by error-prone amplification of the sequence encoding the C-terminal domain of M.HhaI.

## Discussion

The substitution of the catalytic Cys in monomeric C5-DNMTs by Gly, results in the formation of high affinity nucleoprotein complexes, which under certain conditions have lethal consequences for host cells (Wyszynski et al, 1992; Mi and Roberts 1993). In contrast, when a plasmid encoding the same mutation is expressed as a synthetic dimer, there appears to be only a minor impact on cell viability. However, comparisons of cell growth and plasmid transformation efficiencies, conceal a profound difference in the host response to the expression of these two forms of the same point mutation.

Previous studies of this unusual mutant have shown that dissociation from its recognition sequence is approximately 50-fold slower than wild type enzyme and other point mutations (Mi and Roberts, 1993). A similar effect has also been observed when the catalytic Cys is replaced by Ala (Wyszynski et al 1992). However, Gly and Ala are the only two proteogenic amino acids that share a lack of nucleophilic character and possess a side chain volume smaller than that of Cys. The ProGly mutant meets all of the general criteria for a lethal, gain of function mutant.

In a detailed study of the regulated, low level expression of a Pro-Ala mutant of M.EcoRII, Henderson and Kreuzer (2005) used a wide range of repair deficient mutants to compare responses to these types of nucleoprotein lesion. These results point to the involvement of genetically distinct pathways in the resolution of M.EcoRII lesions compared with those involved in the elimination of similar covalent C5-DNMT chromosomal complexes induced by administration of 5-azacytidine or fluoroquinolone. While G:5-azacytidine base pairs in the genome trap C5-DNMTs, the mode of action of the fluoroquinolones involves replication stalling arising from the irreversible formation of DNA gyrase related chromosomal complexes prior to post-replication segregation (Bax et al, 2019). This suggests that the cell marshals subgroups of genes in a combinatorial manner to mount a response to the various types of nucleoprotein complexes that threaten normal genetic transactions.

The valency of macromolecular ligand:protein interactions has been widely discussed in the context of immunoglobulins (Karush, 1976) and considered from a thermodynamic perspective by Mammen et al (1998) and Errington et al (2019). In considering the different outcomes of the binding of monovalent and bivalent polypeptides to specific sequences of DNA, the net fraction of a given concentration of protein bound to a site is simply a function of the number of specific base pairs in the recognition site (a 4 bp sequence occurs on average once every 256 base pairs (44) etc) and the distance between the two recognition sites in three-dimensional space (see Saiz and Vilar 2006 and Laurens et al, 2012 for a discussion of these issues). In one example, the vertebrate CTCF DNA binding protein creates an extensive three-dimensional network of interactions through multivalent binding via its zinc finger domains (Heath et al, 2008; Wang et al, 2019). In prokaryotes, the topological status of the chromosome is also known to be a major factor in genome maintenance, repair and several genetic transactions (reviewed in Uphoff and Sherratt, 2017). Against this background, it is perhaps unsurprising that a multivalent, high affinity DNA binding protein, such as a GST-C5-DNMT ProGly mutant, elicits a different intensity of “DNA repair”, compared with a monovalent form of the same protein.

Most missense mutations have little or no impact on the function of a protein coding gene, unless the base change introduces a stop codon or replaces a critical active site or structural amino acid. The resultant reduction in activity associated with most mutant proteins is only partial, and rarely presents a life or death outcome for the cell. Moreover, where a gene encodes an oligomeric protein, such as a homo-tetramer (α4), the same is usually the case, unless subunit interactions play a key role in the regulatory function of that protein. However, as demonstrated here, the substitution of the catalytic Cys by an amino acid of shorter chain length, and lacking nucleophilic character, has a major impact on whole cell physiology. The results presented here point to a profound difference in the host cell response to a lethal gain of function point mutant resulting from differences in the oligomeric state of the mutant enzyme. In this context, all prokaryotic C5-DNMTs are monomeric apart from a small number of enzymes where their active site is constructed from two polypeptide chains (which has been mimicked in this work with the split domain, b-ZIP constructs). *In vivo*, bacterial C5-DNMTs catalyse sequential methylation reactions, mostly on hemi-methylated, newly replicated genomic DNA, unlike the Type II restriction endonucleases which cleave two palindromic half sites and are almost always closely coupled homodimers.

A series of events, consistent with the results presented here is proposed in Fig. 8. Following transformation of a host cell with a plasmid encoding a monovalent ProGly mutant, transcription and translation generate a low level of the ProGly monomer. As the concentration of ProGly mutant rises, high affinity nucleoprotein lesions will accumulate on plasmid and genomic DNA, where there are several thousand potential recognition sites. It is well established that both plasmid and genomic DNA is a substrate for restriction and modification enzymes. Failure to recover from this state, under selective pressure from an applied antibiotic, is manifest by extremely poor transformation efficiencies and by growth inhibition (Wyszynski et al, 1992; Mi and Roberts 1993). Henderson and Kreuzer (2005) have shown that this observed “toxicity” arises through a major block to replication. In contrast, the negative impact of multivalent ProGly mutants (eg GST-HhaIC81G) has only a minor impact on transformation efficiency. One explanation for the results observed with multivalent C5-DNMTProGly mutants is shown in Fig. 8. Expression of a multivalent C5-DNMT mutant leads to the formation of both *cis* and *trans* nucleoprotein complexes. Multivalent ProGly mutant proteins will make monovalent-like interactions with both plasmid and chromosomal DNA; they will also make a second interaction which, if the site is on the same plasmid molecule or within the same chromosome, can be considered as intramolecular, and will contribute to a net reduction in the dissociation constant associated with multivalent compared with monovalent nucleoprotein complex formation (see Zhou, 2006 for a discussion of valency in macromolecular interactions).

**Fig. 8.**
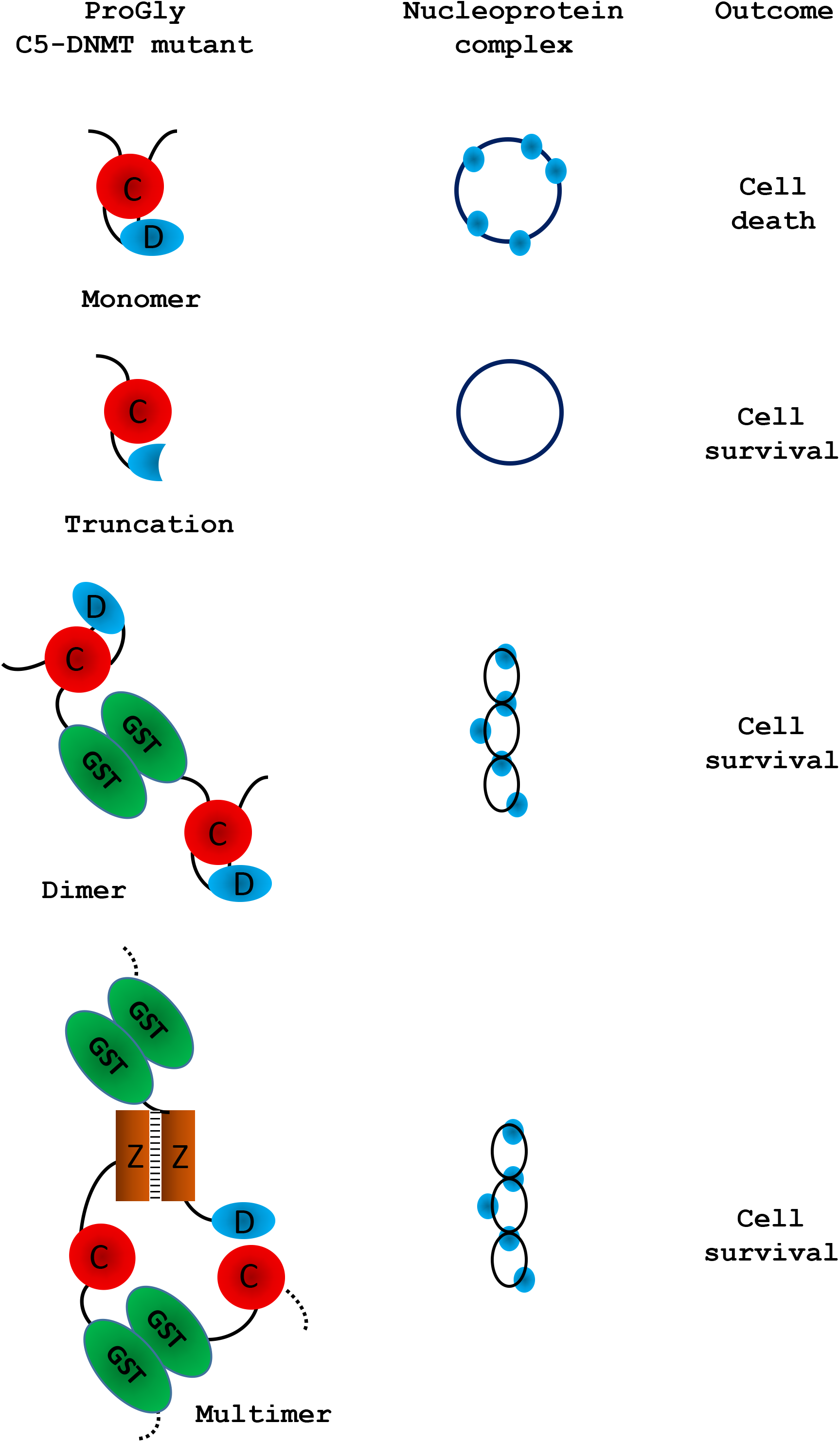
Schematic representation of the observed effect of expression of various forms of the ProGly mutants following transformation of cells. Only multivalent complexes elicit the mutational rescue required for cell survival: these are illustrated by a twisted circle: the ProGly mutants are shown as simple blue circles when bound to DNA.

One of the most striking observations in these experiments is the substantial levels of structural rearrangement observed in those plasmids encoding multivalent ProGly DNMT mutants. Moreover, colonies recovered from a single petri dish typically contain several plasmid variants: multiple experiments revealed similar phenomena, and in each experiment different “structural” classes of mutants emerged following each transformation (inferred from restriction mapping). Finally, the nucleotide sequences of over 100 recovered plasmids were obtained, providing evidence for a general mutational response, removing the need to invoke a specialised mechanism to explain the results presented here. Evidence that the gain of function associated with the ProGly mutation is associated with irreversibly base flipping is strongly supported by the results of a secondary mutagenesis experiment. Mutational rescue from a bivalent ProGly M.HhaI mutant can be mitigated by secondary point mutations that map onto residues associated with DNA recognition and stabilisation of the flipped target cytosine as shown schematically in Fig. 7. In the original structure determination of M.HhaI complexed with a cognate DNA duplex, residues 233-240 and 250-260 were shown to form the interface with the major groove of DNA. The contribution of one specific residue, Gln237 has been the subsequent focus of a detailed structural analysis and is one of the mutants recovered in this work. Daujotyt et al (2004) have presented evidence for a role for this residue in M.HhaI in lowering the energy barrier to eversion of the target cytosine. It seems reasonable to propose that the mutants isolated in this screen neutralize the ProGly phenotype, acting as intragenic suppressors of inappropriate high affinity recognition of DNA which in turn eliminates the need for a mutational rescue event. It is clear from these data that there are many mutagenic strategies available to the host that can overcome the lethality of the ProGly mutant phenotype, but that it is only when the gain of function mutant is expressed in a multivalent context that these solutions are exploited effectively.

In 2001, Loeb described the concept of a mutator phenotype (Loeb, 2001) in which such a cell is characterised by higher than normal levels of mutagenesis, associated with aberrant DNA replication and repair functions. It seems clear that the presence of the ProGly mutant C5-DNMT, when expressed as a bivalent, high affinity DNA binding protein, induces such a phenotype in the host. We suggest that there is a much more rapid (or at least more efficient) induction of error prone repair pathways in response to multivalent compared with monovalent high affinity nucleoprotein complex formation and we refer to this phenomenon as “mutational rescue”. As a result of mutational rescue, transformation efficiencies approach WT levels, and while all surviving cells contain plasmids conferring antibiotic resistance, the GST-C5-DNMT open reading frame carries a variety of inactivating mutations. Interestingly, we observed no evidence for any widespread mutagenesis across the chromosome. Since the formation of ProGly complexes is not expected to be restricted to plasmid sites, the possibility exists that mutagenesis related events that result in the elimination of sequences encoding the toxic multivalent ProGly protein, occur rapidly following plasmid transformation, thereby minimising chromosomal damage., or that replication and subsequent cell division are delayed A more extensive set of genome sequencing would be necessary to test this more thoroughly and may have some relevance to the emerging observations of catastrophic genome re-modelling in the cancer related phenomenon of chromothripsis (Ly and Cleveland (2017).

In the majority of experiments described here, expression of ProGly mutants was achieved following plasmid transformation and analysis was carried out using a number of strategies to optimise transformation efficiency. However, while plasmid analysis provides an insight into the frequency and nature of the mutational rescue phenomenon, it overlooks events on the chromosome. Indeed, following expression of the high affinity C5-DNMT mutant in host cells, nucleoprotein complexes are just as likely to form at genomic sites, since restriction and modification is a well-established genome wide phenomenon and is not restricted to bacteriophage or extra-chromosomal elements. Analysis of the genome sequences of colonies harbouring plasmid indels, revealed no evidence for significant genome wide perturbation. Clearly, non-viable mutants would not have been recovered, but there was no indication of any significant indel events. Fig. 6 shows an alignment of the genomes analysed, with the strain *E.coli* DH5alphaMCR acting as a reference point. The *mcr*- phenotype arises from a deletion of this locus as confirmed by the whole genome sequence analysis. In addition, there was no evidence for anomalous levels of transposition as judged by the alignment of know transposable element sequences.

In this work we have shown that in contrast to the dominant negative phenotype associated with *in vivo* expression of the ProGly mutation in monomeric DNMTs, when the mutants are expressed as synthetic dimers (or higher order oligomers), the lethal effect is abrogated by a raised level of general mutagenesis. While transformants are readily obtained following introduction of the ProGly mutant in artificially oligomerised C5-DNMTs, the input plasmids are often significantly altered and contain a complete spectrum of mutations ranging from missense mutations, to nonsense mutations and a variety of indels. Using several different C5-DNMTs, recognising different DNA sequences and by engineering in multivalency through different oligomerisation mechanisms; the efficient recovery of deactivated ProGly encoding plasmids was shown to be a robust phenomenon.

Application of the fundamental principles underpinning Molecular Biology is currently being “stress tested” by Systems Biologists which in turn will improve outcomes in the field of Synthetic Biology. The successful delivery of programmable synthetic organisms and novel synthetic macromolecules draws heavily on the original model of bacterial gene regulation proposed by Jacob and Monod (1961), and the subsequent validation work on the *lac* operon and the gene regulatory networks in bacteriophage Lambda. In a more recent study, Chure et al (2019) present a systems-based mathematical model based on the relationship betweenthe specificity and interaction affinities of a series of Lac repressor mutants. These kinds of experiments will undoubtedly provide the basis of a more robust framework for designing controllable microbial systems. As the authors point out, while the impact of point mutations on individual genes has provides us with considerable insight into their function, obtaining a wider systems level perspective is now technologically possible and will undoubtedly strengthen future strategies for incorporating programmable functions into cells via genome engineering. The conversion of wild type C5-DNMTs into ultra-high affinity DNA binding proteins, provides a novel platform for investigating the cellular responses arising from interference with genomic transactions. In contrast, from an evolutionary perspective, the benefits of the intrinsic bivalency of immunoglobulins provides the most compelling example of how valency can impart a positive impact on biological function. In the series of experiments, the naturally homo-dimeric GST moiety provides a simple means of tethering together two otherwise independent DNA recognition elements. Assuming equilibrium binding to DNA remains below a certain threshold, both catalytically active and inactive DNMT proteins, monovalent of multivalent, do not threaten cell survival. However, beyond this threshold, systems are in place in *E.coli* that allow cells to respond to the threat from multivalent high affinity DNA recognition far more efficiently than the monovalent threat. This finding could also provide a novel approach to investigating genes involved in the coordination of DNA repair and chromosome packaging in maintenance of genome integrity.

## Experimental Section

### 1. Construction of Mutants

The genes encoding M.*Msp*I, M.*Hha*I (a kind gift from Dr. R. J. Roberts, New England Biolabs, USA) and M.SPRI (a generous gift from Prof. T.A. Trautner, Berlin, Germany) were amplified via the polymerase chain reaction (PCR) using primers to facilitate subsequent ligation of the PCR products into the glutathione-S-transferase encoding vectors (Life Technologies). The M.*Msp*I PCR product was initially inserted into the *Bam*HI and *Eco*RI sites in pGEX 2T, the M.*Hha*I PCR product was ligated into the *Eco*RI and *Xho*I sites in pGEX 4T whilst the M.SPRI gene was inserted into the *Bam*HI and *Xho*I sites in pGEX KG. The genes encoding the above of C5-cytosine-specific DNA methyltransferases (DNMTs) (see Table 1) were constructed using standard protocols (as described earlier (Taylor et al, 1993). All C5-DNMTgenes expressed from pGEX vectors are dimeric, whilst all other constructs yield monomeric (the typical form) enzymes. The details of all constructs are available online at (to be deposited)

In order to introduce the apparently “toxic” ProGly mutation, a number of strategies were employed. In one approach, following the introduction of a pair of unique restriction sites flanking sequence encoding the catalytic Cys residue in M.MspI (Fig.2a), insertion of an oligonucleotide duplex facilitated the substation to create a ProGly mutant. The details are as follows: an engineered *Bsp*EI site was added 30bp upstream of the Pro Cys codons and an engineered *Nde*I site, 5 bp downstream of the Pro Cys codons as shown in Fig. 2a. Alternatively, the C5-MTase was first inactivated by insertion of a kanamycin cassette into the *Xba*I site of the M.*Msp*I gene (or similarly into the *Sal*I site of the M.SPRI gene or an engineered *Xba*I site in the M.*Hha*I gene). Finally, the catalytic and DNA recognition domains were expressed from separate compatible plasmids shown schematically for M.SPRI in Fig 4a. The N-terminal sequence up to the boundary of the Target Recognition element was appended via PCR with a sequence encoding the bZIP sequence from C/EBPα, [LELTSDNDRLRKRVEQLSRELDTLRGIFRQL] and inserted into pACYC184. The sequence encoding the C-terminal domain of M.SPRI was first appended with an N-terminal bZIP sequence and then inserted downstream of the GST open reading frame in the vector pGEXKG, creating an in-frame fusion as shown schematically in Fig. 4. pNSPRIGly-ZIP was maintained in *E.coli* AB1157 cell, which were then made competent prior to a second transformation by pCSPRI-ZIP.

Plasmids were used to transform the various strains of *E.coli* following ligation of the appropriate mutant constructs (Fig.2b). Transformation efficiencies of all strains were monitored and normalised using a stock solution of pLITMUS28 (New England Biolabs, USA). In addition, pGEX 2T encoding a frameshifted M.SPRI mutant (pSPRIGly-2-Stop) was used to determine the efficiency of transformation of each strain. The properties of each plasmid employed in this investigation are listed in Table I and all novel plasmids were characterised by nucleotide sequencing either using the Krebs Institute Biomolecular Synthesis laboratory or the University of Sheffield Medical School DNA sequencing facility (using an Applied Biosystems Model 373A sequencer).

### 2. Transformation, plasmid and genome analysis

All transformations were carried out using strains prepared using the rubidium chloride method described in (Taylor et al, 1993) and transformation efficiencies were routinely evaluated using a standard batch of the commercial supercoiled plasmid pLITMUS38 (NEB). All *E.coli* strains used were obtained from New England Biolabs, except for *E.coli* AB1157 which was a kind gift from Barry Hall (currently Bellingham Research Institute). Plasmids were recovered using mini-prep kits from Qiagen. Restriction analysis, plasmid nucleotide sequencing and DNA methylation analysis was carried out as described elsewhere (Taylor et al, 1993) and genome sequencing was carried out either by the genome sequencing services at Queen Mary University or the University of Birmingham.

## Acknowledgements

This work was supported by studentships awards as follows: PJH (BBSRC), AMA, AAABD, AAA-G, SA-H (Government Scholarships, Saudi Arabia), LA, AA (Government Scholarships, Libya), MA-S (Government Scholarship, Iraq). The authors are grateful for many discussions with Professor Tomas Lindahl (The Francis Crick Institute), Dr. Clive Price (University of Lancaster), Professor Stephen Halford (The University of Bristol) and Professor Barry Hall (currently at The Bellingham Research Institute).

